# Fast *and* long-term super-resolution imaging of ER nano-structural dynamics in living cells using a neural network

**DOI:** 10.1101/2024.07.30.605742

**Authors:** Johanna V. Rahm, Ashwin Balakrishnan, Maren Wehrheim, Alexandra Kaminer, Marius Glogger, Laurell F. Kessler, Matthias Kaschube, Hans-Dieter Barth, Mike Heilemann

## Abstract

Stimulated emission depletion (STED) microscopy is a super-resolution technique that surpasses the diffraction limit and has contributed to the study of dynamic processes in living cells. However, high laser intensities induce fluorophore photobleaching and sample phototoxicity, limiting the number of fluorescence images obtainable from a living cell. Here, we address these challenges by using ultra-low irradiation intensities and a neural network for image restoration, enabling extensive imaging of single living cells. The endoplasmic reticulum (ER) was chosen as the target structure due to its dynamic nature over short and long timescales. The reduced irradiation intensity combined with denoising permitted continuous ER dynamics observation in living cells for up to 7 hours with a temporal resolution of seconds. This allowed for quantitative analysis of ER structural features over short (seconds) and long (hours) timescales within the same cell, and enabled fast 3D live-cell STED microscopy. Overall, the combination of ultra-low irradiation with image restoration enables comprehensive analysis of organelle dynamics over extended periods in living cells.

## Introduction

Super-resolution microscopy (SRM) has enabled visualizing cellular structures beyond the diffraction limit and has been established as an important tool in cell biology research ^[1]^. However, static structures in a cell only reveal part of the picture as biomolecules and cellular structures in a cell are highly dynamic. Understanding these dynamics is therefore crucial for attaining a comprehensive mechanistic picture. Stimulated emission depletion (STED) microscopy is one SRM method that can be employed in living cells and can theoretically achieve unlimited spatial resolution ^[2]^. This resolution is achieved by employing high excitation laser powers (∼kW/cm^2^) and very high depletion laser powers (up to ∼MW/cm^2^), which leads to photobleaching and phototoxicity ^[3]^. In addition, the acquired data could be artifact-ridden as a high laser illumination might induce cellular processes that influence biomolecules in a cell. Various strategies have been introduced to mitigate both photobleaching and phototoxicity. Two simple methods involve engineering highly photostable organic fluorophores ^[4]^ or using transient, weak-affinity and exchangeable labels or fluorophores ^[5–7]^. In the case of photostable fluorophores, photobleaching can only be relieved in the short-term, whereas in the case of transient labels photobleaching can be circumvented relatively longer due to the exchange of fluorophores being sampled. In both cases though, laser powers remain more or less the same and hence, phototoxicity is still a hurdle in the long-term (∼ several minutes). Techniques have been developed that involve minimal sampling which is either controlled by the presence of signal ^[8,9]^ or by the initiation of an event ^[10]^ thereby reducing the overall illumination laser intensity in a cell. However, over the long-term of several hours, phototoxicity still remains. One method that might be promising on this aspect is content-restoration (denoising) of images recorded with low laser intensities (resulting in noisy images), which can be achieved with neural networks ^[11–14]^. Although the maximum time acquisition shown in the case of STED has been limited to several minutes ^[14]^, the technique should enable long-term imaging over several hours due to the decrease in laser intensity incident on the sample.

An additional concern, especially in live cells, is the speed of imaging which determines the accessible temporal dynamics that can be observed in live cells. Focussing on organelle dynamics, the optimum imaging condition would be the one that gives the best possible resolution without any movement artifacts. Depending on the organelle, this could mean that STED nanoscopy yields only sub-par resolution with decreased signal. One such organelle that is highly dynamic in an eukaryotic cell is the endoplasmic reticulum (ER). The ER is the largest organelle in an eukaryotic cell and encompasses different functions ranging from protein folding, translocation and secretion to lipid synthesis and Ca^2+^ signaling ^[15]^. Its structure varies from tubes, sheets, and matrices ^[16,17]^ and has been shown to restructure itself in the order of seconds ^[18]^. To capture the dynamics of the ER, very fast imaging would be required, which in turn means collecting fewer photons from a sample. The integration of a neural network that restores the image content through denoising would mean an increase in the speed of imaging, making it possible to follow organelles like the ER with STED nanoscopy with very minimal to no movement artifacts.

Over the last few years, there have been tremendous improvements in integrating various imaging strategies and their outputs with neural networks for augmented microscopy, image -restoration, -generation, -segmentation, etc ^[11,19–23]^. Two prominent neural networks that have been established for image restoration through denoising and feature extraction via segmentation are UNet-RCAN ^[14]^ and ERnet ^[24]^ respectively. UNet-RCAN employs a two-step architecture for prediction where a U-Net ^[25]^ predicts the broad contextual information, while a residual channel attention network (RCAN) ^[26]^ focuses on the fine details of the image. UNet-RCAN has been shown to outperform commonly used supervised neural networks such as pix2pix ^[12]^ and CARE ^[11]^ in the task of denoising STED images and has been used to reconstruct noisy STED images of fixed and live-cells in both single-plane and volumetric imaging ^[14]^. ERnet, on the other hand, employs a vision transformer-based architecture to classify image pixels as tubes, sheets, sheet-based tubes (SBTs) and background widefield, confocal, and structured illumination microscopy (SIM) images of the ER ^[24]^.

Here, we demonstrate long-term live-cell ER STED imaging using UNet-RCAN for low-intensity denoising and ERnet for segmentation and feature extraction of the ER in the predicted image and establish a robust pipeline for studying live-cell ER dynamics. For this, we trained UNet-RCAN networks using ground truth and low-intensity STED images of live-cell ER. This network, once trained, exhibits the capacity to generate super-resolved images based on their low-intensity counterpart. We report a mean restored resolution of 89 nm in the predicted live-cell images. Whereas live-cell organelle imaging using STED has been limited in mitochondria to 12 min with fluorophores like PKMO ^[27]^ or 12.5 hours (with a snapshot acquisition of one frame every 3.4 minutes) with MAO-N_3_ ^[28]^ and in ER to ∼20 min with a ceramide lipid Cer-HMSIR ^[29]^, we extend this further by acquiring dynamics of ER in resting cells for up to 7 hours in 2D (time resolution: 4.6 s) and up to 17 min in 3D (time resolution: 10.2 s).

To conclude, we show that by combining neural network based denoising to low intensity STED nanoscopy ER dynamics can be observed over the span of several hours under super-resolution and this data can be segmented by tools like ERnet. In addition, our work encompasses the need for incorporating deep learning-based restoration and segmentation strategies to study long-term live-cell organelle dynamics beyond the diffraction limit.

## Results

### Denoising pipeline for live cell ER STED images

We introduce long-term STED microscopy in live cells using low-intensity image recording to minimize photobleaching and phototoxicity. For content restoration through denoising of low-intensity images, the UNet-RCAN network was chosen, as it preserves most of the resolution of the ground truth STED images compared to other methods ^[14]^. UNet-RCAN is a supervised neural network, thereby requiring matching pairs of low-intensity (noisy) and high-intensity (ground truth) images (fig. 1A). The noisy and ground truth image pairs were acquired sequentially with the acquisition switching between lines (line-sequential acquisition) of the noisy and ground truth image to mitigate any spatial offset between the image pairs, which would dampen the quality of predictions. The rapid acquisition of noisy images at exceptionally high speeds (∼300-600 ms per frame) minimizes discernible spatial shifts, while any residual offsets were drift corrected using the NanoJ’s registration method ^[30]^. The acquired noisy and ground truth image pairs were then cropped into smaller patches (128 × 128, 256 × 256 or 304 × 304 pixels) and served as training data for the network (fig. 1B). This has the advantage of generating numerous training instances with only a few measurements. Furthermore, processing the entire recorded image is not possible due to limited RAM size on the GPU. To enhance the network’s resilience to potential variations in samples across measurement days, inputs from at least three measurement days were used for training.

**Figure 1:**
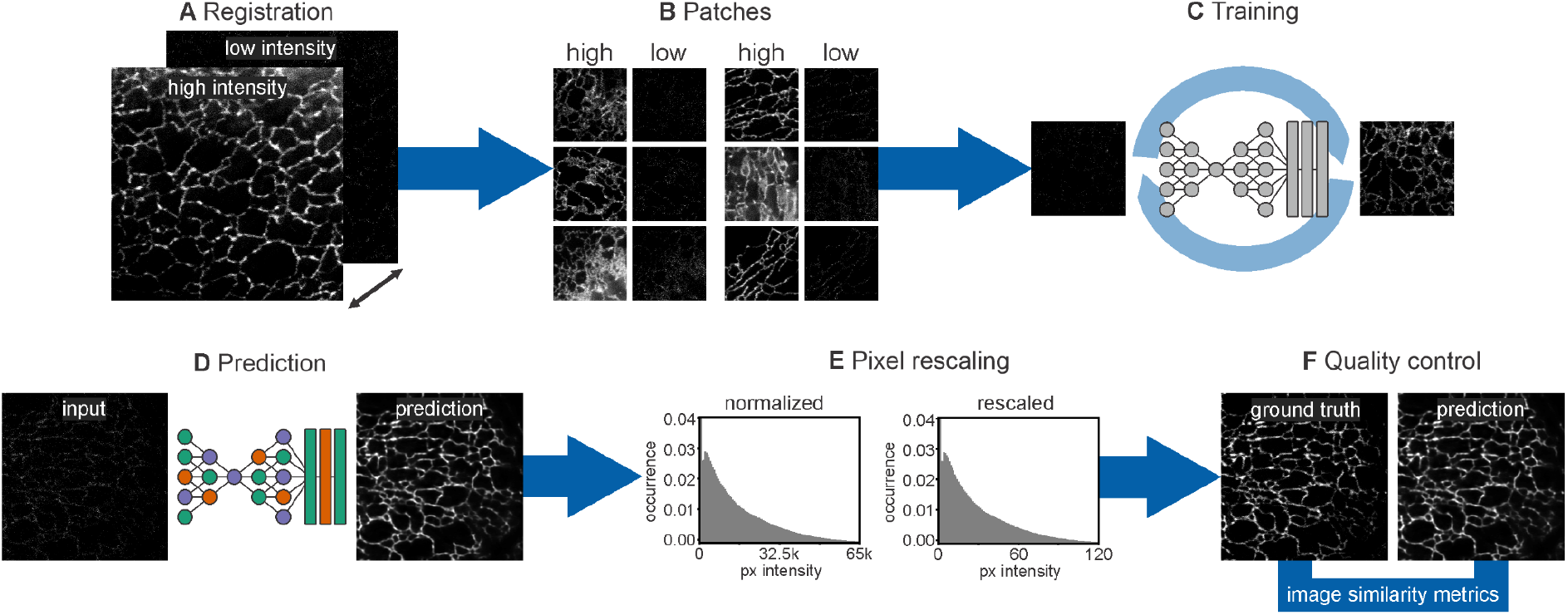
STED denoising workflow. **A** Ground truth and the corresponding noisy images are aligned in the xy plane, **B** from which small patches of high- and low-intensity pairs are generated. **C** A model is generated by training a UNet-RCAN network, wherein the patches from low-intensity images function as input and the high-intensity patches serve as the training target and ground truth. An exploration of various hyperparameters is undertaken to identify a suitable model. **D** The trained model is employed for prediction. **E** The resulting outputs are rescaled to align with the pixel distribution of the ground truth. **F** Quality control metrics are then computed on a designated test dataset to probe the model performance.

Initial attempts to train the UNet-RCAN model using default parameters proved unsuccessful. The model did not converge and a large loss was observed, indicating gradient explosion and rendering the model not useful. Despite adjustments of hyperparameters (patch size, learning rate, kernel init., etc), this issue persisted, necessitating the introduction of additional hyperparameters which were included in the UNet-RCAN code base. An exploration of more than a hundred parameter combinations ensured the identification of a model that converges (SI fig. 1) and performs adequately (fig. 1C).

Model performance was probed with a test dataset, which was acquired on a separate measurement day (fig. 1D). The generated predictions have normalized pixel intensities, which were rescaled to match the pixel distribution of the ground truth data (fig. 1E). The rescaling procedure exclusively leveraged information from the training data, preventing any influence from the test dataset. Finally, quality control metrics were computed on the rescaled predictions and ground truth data, facilitating the identification of an optimal model within the screened parameter space (fig. 1F).

Numerous models were trained with different hyperparameter combinations, allowing for the investigation of their importance regarding model quality with a random forest approach (SI fig. 2). The two most important hyperparameters for model training are an appropriate learning rate and the size of the input. A patch size of 304 × 304 pixels (the size must be divisible by 8 due to the three down-sampling layers in the U-Net) was the optimal choice for this dataset, while larger patch sizes were limited by memory restrictions. Given the pivotal role of this parameter, we advocate for its optimization in conjunction with considerations of image content size, filter size, and downsampling extent. The importance of different parameter values within a parameter group is listed in SI table 1 and 2. This gives an overall indication of appropriate value choices per hyperparameter. However, the best performing models across the hyperparameter search result in slightly different combinations (SI table 3).

### Evaluation of predictions through a comprehensive assessment of image quality

Since the focus of our study was on ER dynamics, we used U2-OS cells stably expressing calreticulin-KDEL as an ER marker conjugated to a HaloTag ^[31]^ which was labeled with covalent HaloTag ligand bound to the fluorophore SiR as a fluorescent system. The live cells were in their normal resting state, which meant that there was minimal perturbation on the ER morphology. The final trained model on the resting cells is termed the resting model. Fig. 2A, B shows ground truth and noisy STED images respectively, of a part of the ER (12 × 12 μm^2^) in a representative cell. The resulting prediction using the resting model recovers the structural details in the ground truth image (Fig. 2C).

**Figure 2:**
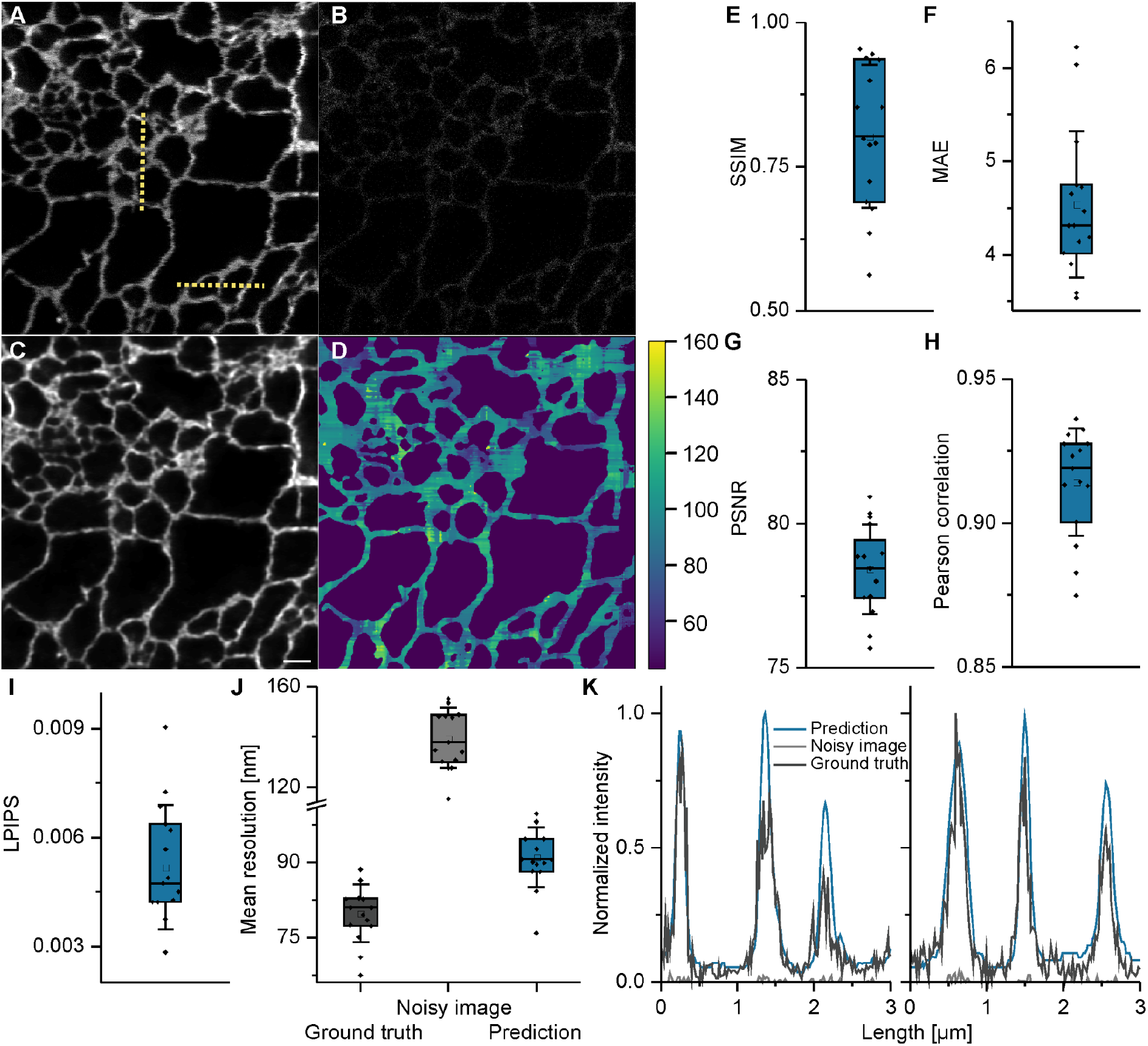
STED denoising of ER in live cells. **A** Ground truth STED image of the ER of a U2-OS cell. **B** Noisy STED image of the ER of a U2-OS cell acquired parallelly as a line acquisition during A. **C** Predicted image of B. using the UNet-RCAN network. **D** rFRC resolution map corresponding to the predicted image, the color code represents the mean resolution in nm. The predicted image quality was probed using different control metrics such as **E** SSIM; **F** MAE; **G** Pearson correlation; **H** PSNR and **I** LPIPS. **J** Mean resolution of ground truth and predicted image calculated using rFRC. **K** Intensity line profiles (along the yellow dotted lines in **A**) of ground truth (dark gray), noisy (gray) and predicted image (blue), the intensities were normalized together as a single group. Scale bar: 1 μm, N = 15.

A rolling Fourier ring correlation (rFRC) map ^[32]^ of the predicted image displays how the resolution varies in different regions of the image giving a mean resolution of 89 nm and a minimum resolution of 42 nm (Fig. 2D). In context, the corresponding ground truth image has a mean resolution of 82 nm. All predicted images were further scrutinized using different quality control (QC) metrics ranging from structural similarity index (SSIM) ^[33]^, mean absolute error (MAE), peak signal-to-noise ratio (PSNR), Pearson correlation to learned perceptual image patch similarity (LPIPS) ^[34]^ (Fig. 2E-I). MAE focuses on the pixel-wise absolute difference between two images. PSNR evaluates the quality of a reconstructed signal, by comparing it to the original, uncompressed signal. The Pearson correlation evaluates the relationship of pixel intensities. Contrary to these metrics, SSIM and LPIPS consider spatial pixel dependencies. SSIM evaluates the similarity of the images in the aspects of structure, contrast, and luminescence. In LPIPS, feature representations extracted from pre-trained neural networks are compared between prediction and ground truth images. This combination of different metrics, along with the spatial resolution, is a robust toolbox for comprehensive image quality assessment. All QC metrics show that the predictions recover the structural details of the ground truth image. In addition, the predictions also recover the resolution although not fully compared to the ground truth images (Fig. 2J). Intensity profiles across the image also show that the predictions retain the underlying details of a ground truth image (Fig. 2K).

### Effects of morphological changes on model robustness

Subsequently, we wanted to explore how the quality of the network prediction is affected if the morphology of the ER undergoes structural changes. For this purpose, cells were treated with Torin1 and Bafilomycin to induce ER-phagy ^[35]^. Noisy ground truth image pairs were acquired from autophagy-induced cells for training a unique model for autophagy induced cells. Autophagy induction is a time dependent process where the ER of a cell changes continuously ^[36,37]^ since induction, therefore we went with a prediction model where the training data was partly from autophagy induced cells and partly from resting cells to account for diverse morphologies. This model was termed mixed model. The mixed model performed well with the test data from autophagy induced cells (SI Fig. 3) and the resting model with the resting cell test data (Fig. 2). This performance extended in terms of structure recovery (SI Fig. 3A-C), resolution (SI Fig. 3D), intensity fluctuations (SI Fig 3E) and QC metrics (SI Fig. 4).

To test the robustness of both the resting and the mixed model we predicted test input data from resting cells with the mixed model and autophagy-induced cells with the resting model and evaluated their performance with QC metrics and resolution. In terms of QC metrics, the mixed model and resting model performed equally well for data of autophagy-induced cells (SI Fig. 4). The model solely trained on resting cells performed slightly better on the resting cell test dataset (SI Fig. 5). Overall, the structural features were restored in all predictions (SI Fig. 6). Interestingly, movement artifacts that partly appear in the ground truth test data due to the fast organelle dynamics are averaged out in the predicted images. This shows that the data from both conditions was within the generalization capability of the resting model

### Strategies to discern structural information from hallucinations

To ensure the quality of predictions, it is crucial that the low-intensity input still contains some signal that has not completely bleached away. In the absence of signal, hallucinations occur (SI fig. 7A and B). To distinguish hallucinated predictions from those stemming from structure, the similarity of adjacent frames in a movie can be probed. Moving organelles exhibit a degree of similarity between adjacent frames, resulting in higher structural similarity when compared to adjacent predictions derived from empty field of views (FOVs) (SI fig. 7C). By tracking the similarity over time the authenticity of predictions can be validated. Areas of uncertain predictions might not only arise temporally, but also spatially; for instance, when a minute signal was collected from out of focus planes, which is not sufficient for a robust prediction (SI fig. 7D-E). By thresholding the pixel intensities of the low-intensity image, the pixels can be roughly grouped into background, uncertain, and signal pixels. This grouping gears the attention to areas with potential artifacts.

### Denoising and segmentation to probe fast and long-term ER dynamics

ER as an organelle exhibits fast dynamics visualized as rearrangement in structure and has been shown to happen in the order of seconds ^[18]^. In addition, the morphology of the ER is also profoundly responsive to conditions that induce stress and is in the order of hours to days. Taken together, it is important to observe ER dynamics with a time resolution of seconds over the span of hours. Therefore, as a first step into this direction, we extended our UNet-RCAN implementation from super-resolved planar images of the ER to study the long-term temporal progression of ER dynamics in resting cells with a time resolution of seconds.

We acquired low-intensity STED images of the ER in normal resting cells for over 7 hours with a temporal resolution of 4.6 s and denoised it using the UNet-RCAN model trained on resting cells (Fig 3A, Video S1). We acquired this data for three different samples seeded over three different days to account for heterogeneity. ER rearrangements are variable in each cell even in their normal resting state (Fig 3A, SI fig. 8x, Video S1-3).

**Figure 3:**
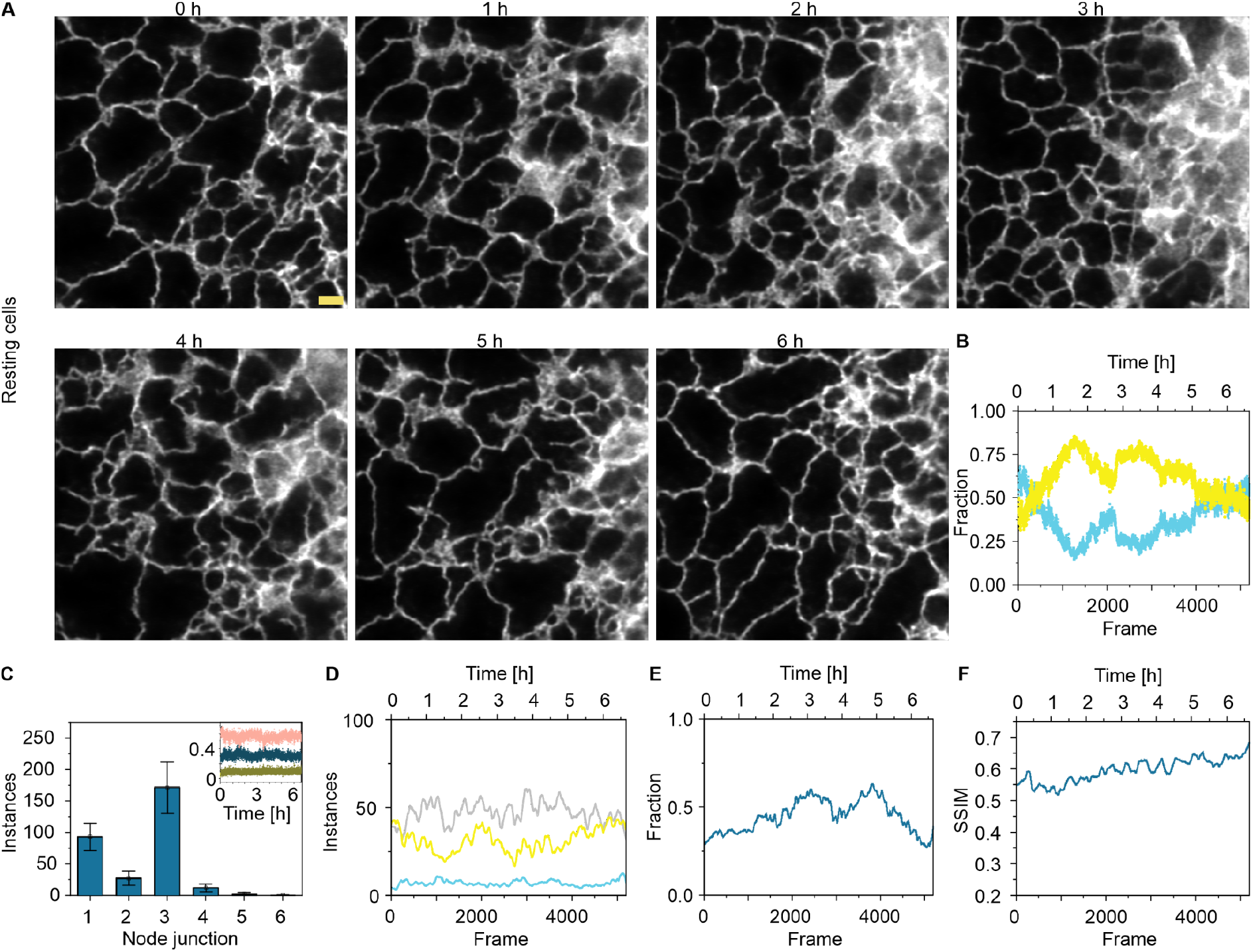
Long term denoising and segmentation of fast ER dynamics. **A** Predictions of low intensity STED images of ER (calreticulin-KDEL) at different time points, acquired on resting cells and predicted using the resting model. **B** Fraction of tubes (cyan) and sheets (yellow) calculated using ERnet segmentation on **A. C** Amount of different tubular junctions calculated using ERnet segmentation averaged over all frames. **C, Inlay** Fraction of one- (green), two- (blue) and three- (pink) way tubular junctions over the whole measurement time of **A. D** Instances of sheets (yellow), tubes (cyan) and gaps (light gray) calculated from **A. E** Fraction of foreground pixels in each frame in **A. F** SSIM scores for adjacent frames calculated for all frames in **A**. N = 3. Scale bar: 1 μm.

We next aimed to quantify ER rearrangements over the full measurement time in more detail. As a start, we employed a neural network based segmentation tool, termed ERnet ^[24]^, on STED nanoscopy datasets. ERnet takes planar images as input and segments them into tubes, sheets, and sheet based tubes (SBTs). The segmented SBTs resembled part of sheets rather than tubular structures, probably due to the lack of real SBTs in the imaged area and the ERnet model being trained to detect SBTs. In our case, the small spots were merged with the segmented areas of sheets (SI fig. 9). We note that we used an ERnet model trained on ER datasets acquired through SIM microscopy. Despite the differences in imaging modalities, the model demonstrated robust performance with our denoised STED data. ERnet segmentation shows that structural rearrangements are variable among cells in resting state. However, in all cases the amount of sheets and tubes fluctuates around a fraction of 0.5 (Fig 3B, SI fig. 10Ai, ii). In addition, we also see that the number of tubular junctions shows a majority at 3 (Fig 3C, SI fig. 10Bi, ii) as shown before ^[24]^ and this feature stays constant over time (Fig 3C inlay, SI fig. 10Ci, ii).

In addition to ERnet based segmentation, we analyzed structural rearrangements through various additional metrics such as instances of structures, the fraction of foreground pixels and SSIM score between adjacent frames. In the case of instances, the sheer number of sheets, tubes and gaps in between tubes and sheets were calculated for each frame in the whole time series datasets. Fig 3D shows that the number of instances of tubes, sheets and the gaps (background enclosed by ER structure) between them stays more or less constant throughout the imaging time. In the other two cells recorded we see more fluctuations in the case of gaps between tubes and sheets (SI fig. 10Di, ii). The fraction of foreground pixels shows a trend similar to what we observe in the ERnet segmentation and is variable for the three different cells (Fig 3E, SI fig. 10Ei, ii). Here we define foreground pixels as any non-zero pixel in a binarized image of every frame. The SSIM scores reflect the frequency of changes happening in-between frames. Fig 3F shows a slight increase in SSIM through the progression of the image acquisition. Data from other cells show heterogeneity with one staying constant throughout (SI fig. 10Fi) and one showing a slight decrease (SI fig. 10Fii) but overall SSIM values do not fluctuate drastically, reflecting that the frequency of structural rearrangement in resting cells is more or less constant.

### Extending denoising to visualize long-term ER dynamics in 3D

The ER is a highly convoluted structure extending in 3D ^[16]^. Although 2D planar imaging over time gives insights into ER dynamics it does not completely reflect the underlying ground truth. Possible artifacts may arise from 2D projection of the dense 3D ER structure. Therefore, we extended the UNet-RCAN model trained on 2D planar images (resting model) to denoise volumetric images from living cells over time.

We first tested the robustness of the resting model on singular volumetric datasets. The 3D stacks were limited to a thickness of 700 nm and only noisy images were acquired to keep spatial offset between frames to a minimum. In addition, a top-hat point spread function (PSF) pattern was used for the depletion laser to gain axial resolution. SI Fig 11 shows a predicted 3D stack of an exemplary cell visualized as xy, yz and xz planes (Video S4). Since only noisy images were acquired in the 3D stack, we measured additional single plane noisy and ground truth images (SI Fig 12) with the same settings including a top-hat depletion laser PSF as a control. We performed QC metrics on the control dataset (SI Fig 13) to show that the resting model does indeed denoise images acquired with a top-hat depletion laser PSF achieving a mean resolution of 89 nm (SI Fig 13F).

We then used the resting model to predict time series data of 3D stacks acquired over 17 min with a temporal resolution of 10.2 s. Fig 4A shows orthogonal slices of an exemplary predicted data from 3D stacks over different points of time in the acquisition. The 3D surface rendering of the time series data clearly shows that considerable dynamics would be missed, e.g. through projection, if it were not recorded in 3D mode (Fig 4B, Video S5). Fig 4C highlights this point by visualizing certain structural rearrangements over time for one of the frames of the whole 3D stack. Similar data was acquired for different resting cells over different days (Video S6,7). Hence we show that the resting model could be used on z-stacks and extended to 3D time series over 17 min at super-resolution.

**Figure 4:**
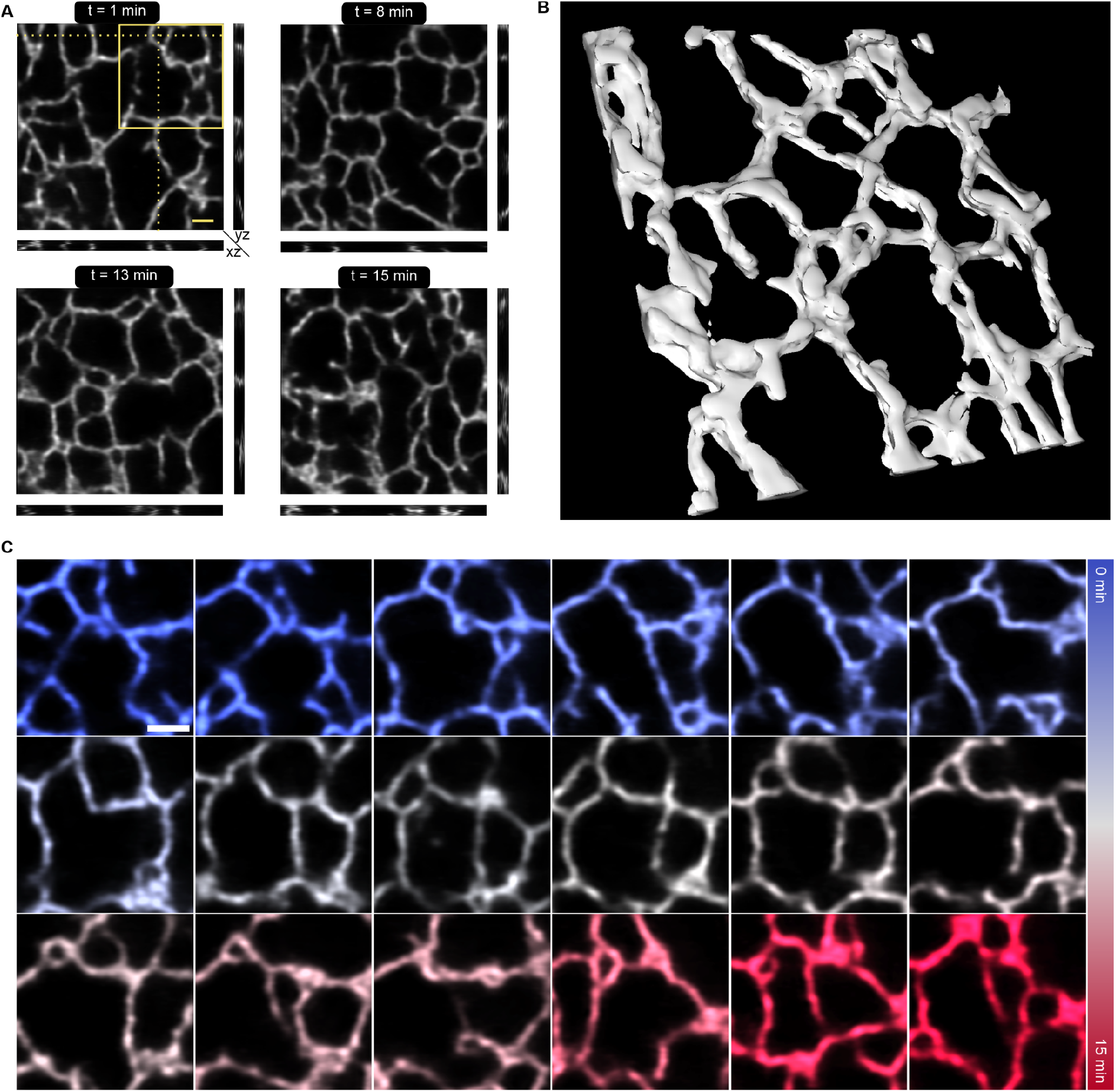
Denoising 3D stacks over time to visualize fast 3D ER dynamics. A Orthogonal projections of predictions from z-stacks of the ER acquired on live-cells at different time points. **B** Surface rendering of one of the z-stacks shown in **A. C** Representative images from the inlay and the same z-plane shown in **A** at the different time points throughout the acquisition. N = 4. Scale bars: 1 μm.

To conclude, we implemented an UNet-RCAN network to denoise STED images (planar- and volumetric imaging) of ER (calreticulin-KDEL) in living cells and by reducing the laser intensity incident on the cells extend STED nanoscopy on live-cells to over 7 hours.

## Discussion

Visualizing organelle dynamics in live-cells with sub-diffraction resolution and for extended time spans so far required either specialized microscopy equipment (lattice light sheet or grazing incidence SIM) ^[18,38]^ or intermittent sampling due to the prevalence of photobleaching and phototoxicity ^[28]^. An alternative approach is to restore image content using a neural network ^[11,13]^. Previous work demonstrated that neural network assisted denoising can restore STED images and mitigate photobleaching and photodamage by reducing the pixel dwell time and hence light exposure. In addition, they showed that their UNet-RCAN network outperformed pre-available networks ^[14]^. The aim of our work was to expand live-cell imaging to the time scale of hours and at the same time with high temporal resolution to visualize ER dynamics. By implementing an UNet-RCAN network, we were able to measure live-cells with ∼70 times lesser light dose per pixel while maintaining a sub-100 nm resolution. We show that a model solely trained with resting cells is robustly predicting both data acquired from autophagy-induced cells and normal resting cells respectively (SI Fig 4 and SI Fig 5). This shows that models generalize morphological changes to some extent, and acquiring training data from resting cells could be sufficient when working with multiple conditions. By leveraging the ∼70 times reduction in light dose and a faster acquisition speed of 600 ms per frame, we were able to measure low-intensity STED images of ER (calreticulin-KDEL) in living cells for over 7 hours with minimal movement artifacts.

We then predicted super-resolution STED images from these low-intensity images giving rise to time series data with sub-100 nm spatial resolution. The predicted images had a ∼4 s time resolution for a time span of ∼7 hours giving full structural information on the rearrangement of the ER. We accessed this by using ERnet based segmentation which is a neural network based semantic segmentation tool that has been shown to classify ER structures into tubes, sheets and SBTs especially from data acquired through SIM microscopy ^[24]^. In addition, through ERnet segmentation, Lu et al., have shown that three way tubular junctions are dominant in healthy ER and they observed that in 20% of the healthy cells that they analyzed exhibited ER with higher order tubular junctions ^[24]^. Both of these conditions are similar to what we observe in our time series data, where the three way tubular junctions dominate and higher order junctions are also evident (Fig 3C, SI fig 10B). We also show that structural rearrangements could be analyzed through other metrics such as the foreground pixels and the SSIM index. Whereas the foreground pixel information is similar to that extracted from the ERnet, though not as detailed, the SSIM index gives information on the frequency of rearrangements occurring between frames. We see that for two of the three cells measured, the SSIM index stays constant throughout the measurement indicating that the structural rearrangements happening are not fluctuating from the mean. In the case of the outlier, we can see from Video S3 that the cell itself moves and the amount of ER in the FOV is decreased which could lead to the decrease in the SSIM index. In addition, we also used the resting model for predicting volumetric data acquired from living cells. It can be seen from the control test data (SI fig. 13F) that the mean resolution achieved is 89 nm. We extended this further by applying it to volumetric time series data for up to 17 min. We can see that certain structural rearrangements could not be comprehended if it were not for volumetric imaging (Fig 4C, Video 5-7). Volumetric imaging might prove useful especially when studying inter-organelle dynamics. Taken together, the resting model we trained with the UNet-RCAN network is robust enough to predict not only 2D planar images but also 3D volumetric images and 2D planar images from stress induced cells.

We chose the ER as the target organelle due to its dynamics being largely unknown. The morphology of the ER is profoundly responsive to conditions that induce stress within a cell. In this context, the ER plays an important role in stress-induced autophagy, a cellular process, which involves the degradation and recycling of cellular components, defined as ER-phagy. Only fixed cell studies have been performed on ER-phagy samples under super-resolution to the best of our knowledge ^[35,39]^. Our pipeline that we developed here would serve in the future to visualize fast and long term ER dynamics under different stress-induced conditions with different cell line backgrounds. Our work mainly focused on the ER which was recorded in a single fluorescence channel. We envision extending this approach to multiple spectral channels or add additional measurement parameters to study inter-organelle dynamics or intra-organelle dynamics by selecting multiple targets of interest. Future work might include the extension of describing other cellular structures along the ER, drawing a more holistic understanding of organelle dynamics and interactions. We also note that content restoration and denoising are areas of active research in the field of neural network assisted bioimage analysis, and we anticipate that alternative approaches can be integrated into this workflow ^[20,40]^.

In summary, our work demonstrates a neural network based workflow accessing the fast ER dynamics with long time observation at high resolution in planar and volumetric image acquisitions. This robust framework could be further applied to different scenarios, where fast organelle dynamics with sub-100 nm resolution are of interest.

## Methods

### Cell culture

Live-cell samples were prepared from U2-OS cells stably expressing Calreticulin-KDEL-HaloTag7. The cell line was cultured in DMEM/F12 media supplemented with 10% fetal bovine serum (Corning, USA), 1% (v/v) penicillin-streptomycin (Thermo Fisher Scientific, Germany) and 5% (v/v) Glutamax (Thermo Fisher Scientific, Germany). The cells were incubated at 37 °C and 5% CO_2_ and were passaged every 2-3 days or when they reached 80% confluency.

### Sample preparation for live-cell microscopy

U2-OS cells stably transfected with calreticulin-KDEL-HaloTag7 were seeded onto 8-well chambered coverglass (Sarstedt, Germany) at an amount of 1 × 10^4^ cells per well. The cells were incubated overnight at 37 °C and 5% CO_2_ and were induced the next day with 250-500 μg/mL of doxycycline (Sigma Aldrich, Germany) in culture media. Two days post-induction, the cells were incubated for 15-20 min with 300 nM of HaloTag ligand conjugated to the fluorophore SiR diluted in culture media. The cells were then washed thrice with culture media with 5 min incubation at 37 °C with 5% CO_2_ between every wash. The cells were incubated in culture media at 37 °C with 5% CO_2_ till they were taken out for imaging. For imaging, the culture media was then exchanged for live-cell media (reduced culture media with HEPES, Thermo Fisher Scientific, Germany) and the cells were imaged within the next two hours unless explicitly mentioned.

### Autophagy induction

For autophagy induction, the media of the seeded cells were exchanged with live-cell media containing final concentrations of 250 nM Torin1 (Bio-techne, USA) and 250 nM BafA1 (Cayman Chemical Corporation, USA). The cells were imaged immediately to observe the progression of autophagy over time.

### STED microscopy

An Abberior expert line microscope (Abberior Instruments, Germany) was used for performing confocal and STED imaging. The setup had a Olympus IX83 body (Olympus Deutschland GmbH, Germany) where the imaging was done using a UPLXAPO 60x NA 1.42 oil immersion objective (Olympus Deutschland GmbH, Germany). Sample images were acquired by exciting with a 640 nm excitation laser (both confocal and STED imaging) and depleting (in the case of STED images) with a 775 nm laser which either had a donut PSF (for planar and long-term imaging) or a top-hat PSF (for volumetric imaging and planar control) and with a delay of 750 ps and fluorescence photons between 750 ps and 8.75 ns were detected between every laser pulse. Fluorescence was collected in the spectral range of 650 nm to 760 nm using an avalanche photo diode (APD). The power levels of the lasers used and the parameters used for imaging are summarized in Table 1. Note that the power levels for the lasers were measured at the back focal plane. Training data was imaged in a 7.5×10 μm^2^ and 12×12 μm^2^ field of view (FOV); test data was imaged in a FOV of 12×12 μm^2^; 2D time series data was imaged in either 12×12 μm^2^ or 15×15 μm^2^ FOVs and 3D time series was imaged in a FOV of 10×10 μm^2^.

**Table 1:** Imaging parameters used for confocal and STED imaging.

| Imaging modality and condition | Excitation laser power ($\mu\text{W}$) | Depletion laser power (mW) | Depletion laser PSF pattern | Line accumulation /integration | Pixel dwell time ($\mu\text{s}$) | Pinhole diameter (airy unit (AU)) | Pixel size/ Voxel size (nm) |
| --- | --- | --- | --- | --- | --- | --- | --- |
| Ground truth, single plane | 56.3 | 166 | Donut | 6 | 0.5 | 0.81 | 20 |
| Low intensity, single plane; timeseries | 4.8 | 166 | Donut | 1 | 0.5 | 0.81 | 20 |
| Low intensity, z-stack | 4.8 | 166 | top-hat | 1 | 0.5 | 0.81 | 20 |
| Ground truth, single plane | 56.3 | 166 | top-hat | 6 | 0.5 | 0.81 | 20 |
| Low intensity, single-plane | 4.8 | 166 | top-hat | 1 | 0.5 | 0.81 | 20 |
| Low intensity, z-stack, timeseries | 4.8 | 166 | top-hat | 1 | 0.5 | 0.81 | 35 |

### Image processing

High-intensity and low-intensity images were registered with NanoJ-Core’s registration method ^[30]^. For the resting condition, training images were acquired from 3 measurement days. For the mixed model, 26% of images were added from the treated condition. The same amount of data was used for training to allow comparison between the resting and mixed model performance. Images were cropped into patches of 128 × 128 pixels, 200 × 200 pixels, 256 × 256 pixels, and 304 × 304 pixels to determine which patch size is optimal. A training and validation split of 90%-10% was made. Per condition, 15 test images of size 600×600 px^2^ were acquired at a different measurement day.

### Network training

UNet-RCAN networks ^[14]^ were trained on a NVIDIA RTX 3090 24 GB GPU. As the default values led to exploding gradients, we added some additional hyperparameter options to the published codebase, and the parameter space was explored to find a robustly working model. This included different loss functions (leaky ReLU, tanh), kernel initialization strategies (glorot uniform, lecun uniform, orthogonal), gradient clipping (0.1, 0.01, 0.001), and l2 regularization (0, 0.01, 0.001). Different learning rates were additionally tested during the search (0.001, 0.0001, 0.00001 for the resting and 0.001, 0.0001, 0.0005, 0.00005, 0.00001, 0.000001 for the mixed condition), along with different patch sizes (128 × 128 pixels, 200 × 200 pixels, 256 × 256 pixels, and 304 × 304 pixels). The batch size was chosen as the maximum possible number per patch size before an out-of-memory error occurred on the GPU (patch size: batch size, 128: 36, 200: 16, 256: 10, 304: 6).

A parameter space search was executed to identify possible parameter combinations for models that perform well on the test dataset. To diminish the search time, 1000 data instances were included and training was interrupted after 10 epochs or if the loss exceeded a value of 1000 during training. 147 models were evaluated for the resting condition and 393 models for the mixed condition. Quality control metrics were calculated on a test dataset after every model run to determine the model performance. The metrics SSIM ^[33]^, MAE, and the resolution (determined with decorrelation analysis ^[41]^ in NanoPyx v0.2.2 ^[42]^) were each linearly transformed between 0-1 (0 means worst and 1 means best for all scores) across all model runs per condition. A “total score” was computed by summing the transformed scores and dividing them by three to assess performance across multiple metrics. The top 5 models, based on the highest total score, were trained until convergence on the full dataset with 1764 instances.

For the final calculation of quality control metrics, the pixel intensities of the predicted test images must be rescaled, as the model outputs normalized intensities. A scale and offset were calculated based on the pixel values of predicted training instances and their ground truth images; the test dataset is deliberately excluded from this process. A linear scaling operation was applied to the pixel values of the predicted images. Values above the 99th quantile and below the 1st quantile were clipped. The scale and offset values were determined per model and applied to the test data before calculating quality control metrics. The final model was selected based on the highest total score, with parameters detailed in SI table 3.

### Quality control metrics

The MAE was calculated using the mean absolute difference of pixel values. SSIM ^[33]^ and PSNR were computed using the scikit-image library v0.19.3 ^[43]^. Pearson correlation was determined with SciPy v1.9.3 ^[44]^. The learned perceptual image patch similarIty (LPIPS) ^[34]^ was ascertained with a VGG model. Resolution during model training was determined with decorrelation analysis ^[41]^ in NanoPyx v0.2.2 ^[42]^. Resolution for the test data was determined using rolling Fourier ring correlation v0.2.5 ^[32]^ with a 1/7-threshold and a background intensity of 20.

To discern predictions stemming from FOVs with signal and FOVs containing only noise, the SSIM was calculated between adjacent predicted frames of the resting movie and a movie recorded of only dark noise of the detector (in an empty image plane without sample). To spatially group pixels into the categories background, uncertain, and signal, a sliding window of size 10×10 px^2^ was used to sum the pixel intensities within this window per pixel. Borders of the image were padded with reflected pixels to maintain the input size. A pixel was labeled as background, if the sum was equal or below 5, as uncertain if in between 6 and 25, and as signal if above 25. The window size and thresholds must be adapted per use case.

### Feature importance

The *RandomForestRegressor* from scikit-learn v1.0.2 ^[45]^ was used to determine hyperparameter importance based on the 147 model runs for the resting and 393 model runs for the mixed condition. It ranks the hyperparameters based on their importance in achieving a high “total score”. Furthermore, suitabilities are assigned to the different parameter options within a parameter group.

### Structural analysis of the ER

Rolling ball background subtraction with a radius of 15 px was applied to the images in Fiji ^[46]^ prior to ERnet processing. This prevents high background regions from being falsely classified as ER sheets. The pretrained *20220306_ER_4class_swinir_nch1.pth* ERnet v2.0 model was used in Google Colab ^[24]^ to describe the ER properties in a living cell in the predicted long-time movies for the resting and mixed conditions. The segmentation outputs the structures tubules, sheets, and sheet-based tubules (SBTs). As the latter was not visible in the movies and ERNet just segmented some minor spots as SBTs, these spots were merged into the sheet segmentation.

### Post processing volumetric data

The predictions from volumetric datasets (Video S4-7) were checked for hallucination artifacts as shown in SI fig. 7E and peripheral frames were cropped to avoid them. The cropped 3D stacks were rendered as surfaces using the “3D Viewer” in Fiji with “Display as” set to surface, “Resampling factor” set to 2 and “Threshold” set to 50.

### Statistical analysis

All datasets were probed for normality by applying Shapiro-Wilk tests (α = 0.05) and Mann-Whitney-U tests were used to compare the metrics of ground truth and predicted images. Levels of significance were defined as: p > 0.05 no significant difference (n.s.), p < 0.05 significant difference (*), p < 0.01 very significant difference (**), p < 0.001 highly significant difference (***). All tests were performed in Origin Pro 2018b.

## Data availability

Data (raw and predicted datasets) reported in the manuscript are available on Zenodo (10.5281/zenodo.13120793).

## Acknowledgments

This work was supported by the Deutsche Forschungsgemeinschaft (DFG, German Research Foundation) through SFB 1177 (project-id: 259130777) (M.H.), GRK 2566 (project-id: 414985841) (M.H., M.K.) and INST 161/1020-1 FUGG (M.H.). We thank Martin Weigert, Jun-Yan Zhu and Magnus Petersen for the insightful scientific discussions. We thank Anshu Katri and Volker Dötsch for generously providing Torin1 and Bafilomycin A.

## Author contributions

M.H., A.B. and J.V.R. conceived the study. A.B., A.K., L.K. measured the microscopy datasets. J.V.R. set up the and trained the neural network. H.D. setup the GPU server. J.V.R. and A.B. performed data analysis and wrote the original draft manuscript. A.B., J.V.R., M.W., M.G., H.D., M.K. and M.H. reviewed and edited the final manuscript. M.H. acquired funding and oversaw the project.

