## Supplemental figures and tables for "Fast *and* long-term super-resolution imaging of ER nano-structural dynamics in living cells using a neural network"

### Supplemental Information

Supplemental Figures 1 - 13
Supplemental Tables 1 - 3

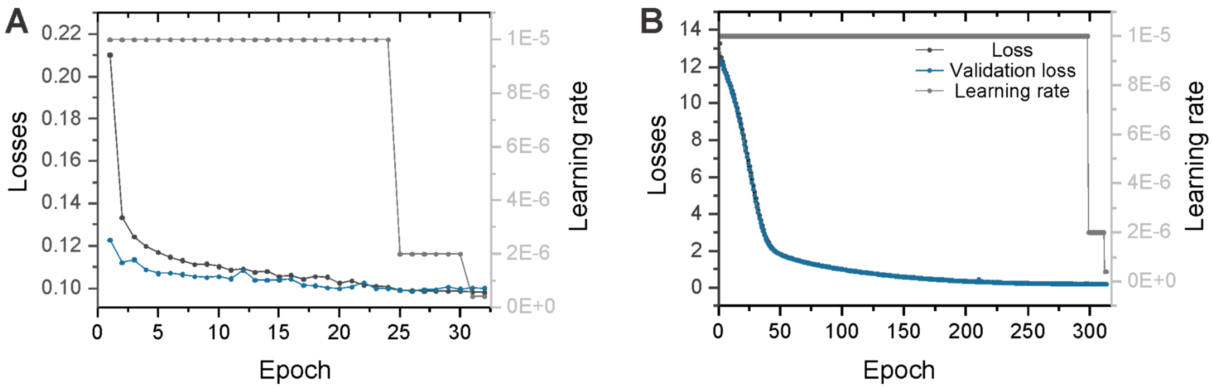

**SI Figure 1: Loss curves and learning rates of the training process.** Training-, validation- loss curves and the learning rates of the **A** resting model and **B** mixed model used in this study. During training, the learning rate was reduced with the LearningRateOnPlateau and the training process eventually stopped if the validation loss did not improve for 6 epochs.

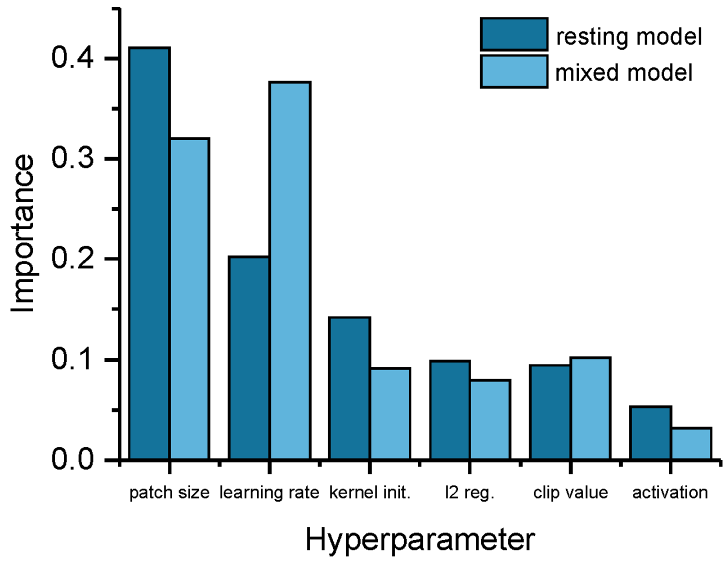

**SI Figure 2: Hyperparameter importance.** For both the models (resting model and mixed model) used in this study, the patch size and learning rate are the most important hyperparameters influencing model performance, followed by kernel initialization, l2 regularization, and gradient clip value in the middle field, and lastly the type of activation function.

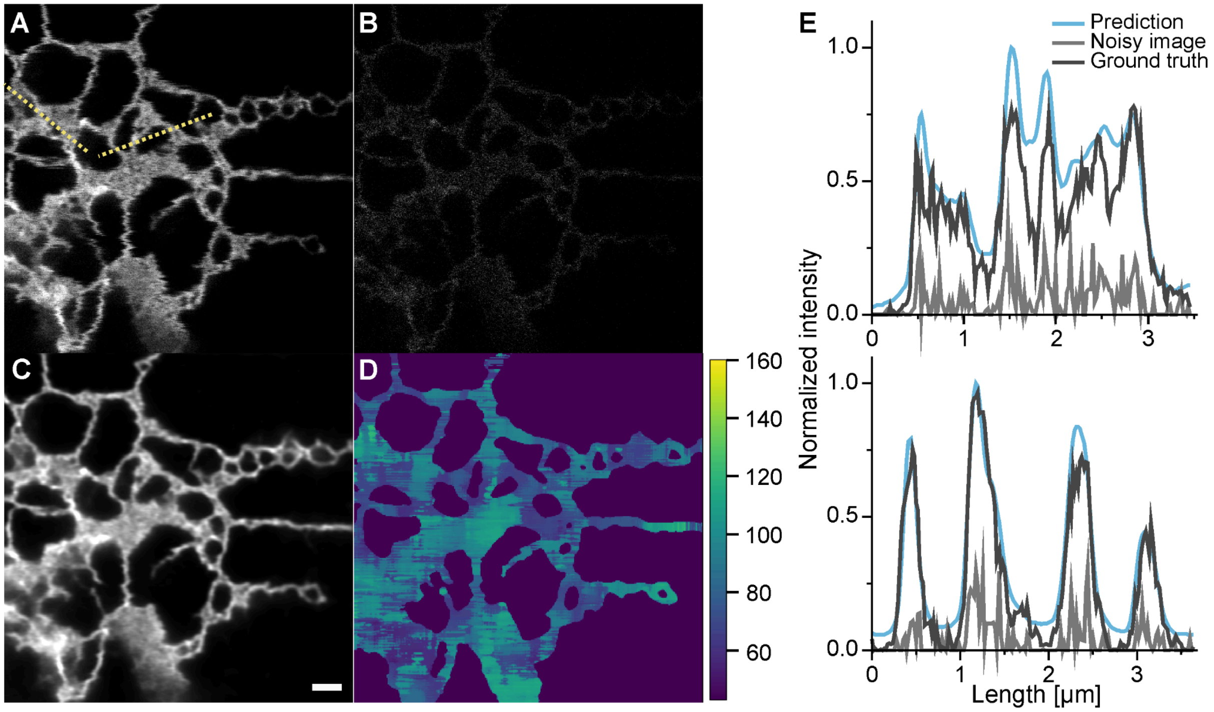

**SI Figure 3:** STED denoising of live cells induced for stress. **A** Representative ground truth and **B** noisy STED image of the ER in a stress-induced live cell and **C** its corresponding prediction. **D** rFRC resolution map of the predicted image pair. **E** Intensity line profiles of ground truth, noisy and predicted image at two highlighted regions of interest. The intensities of all three datasets were normalized together as a single group. Scale bar: 1 µm.

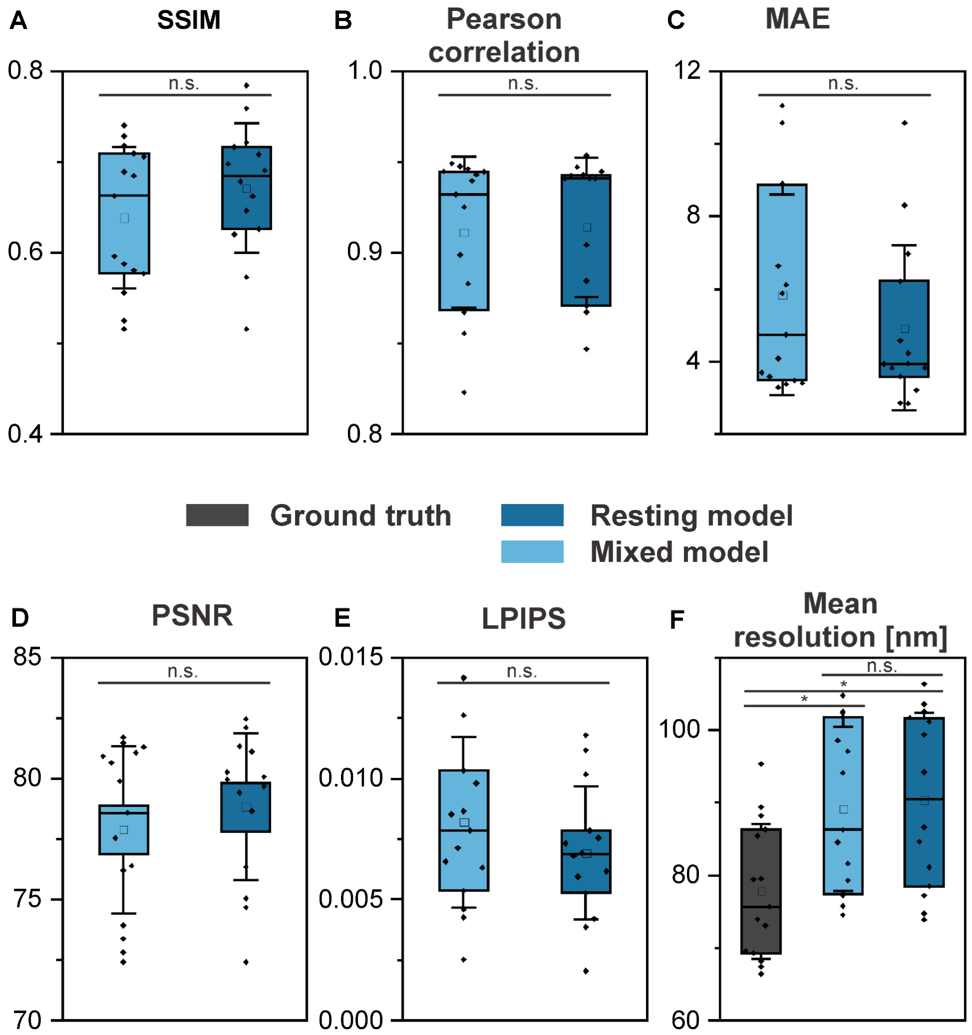

**SI Figure 4**: **Quality control metrics of prediction from autophagy-induced cell dataset.** Quality control metrics **A** SSIM, **B** Pearson correlation, **C** MAE, **D** PSNR, **E** LPIPSand **F** mean resolution from rFRC of datasets acquired from stress-induced cells predicted using either the mixed model (light blue) or the resting model (blue), dark gray: mean resolution of ground truth image, N = 15.

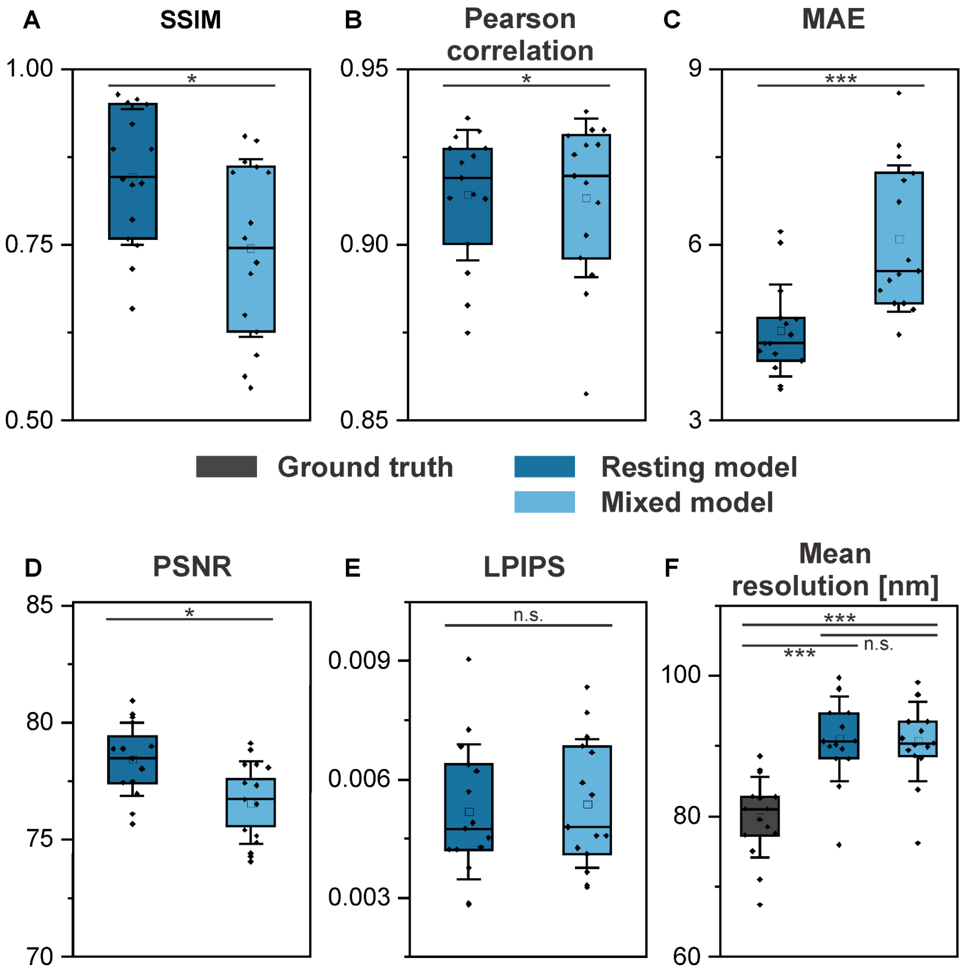

**SI Figure 5**: **Quality control metrics of prediction from the resting cell dataset.** Quality control metrics **A** SSIM, **B** Pearson correlation, **C** MAE, **D** PSNR, **E** LPIPS, and **F** mean resolution from rFRC of datasets acquired from resting cells predicted using either the resting model (blue) or the mixed model (light blue), dark gray: mean resolution of ground truth image, N = 15.

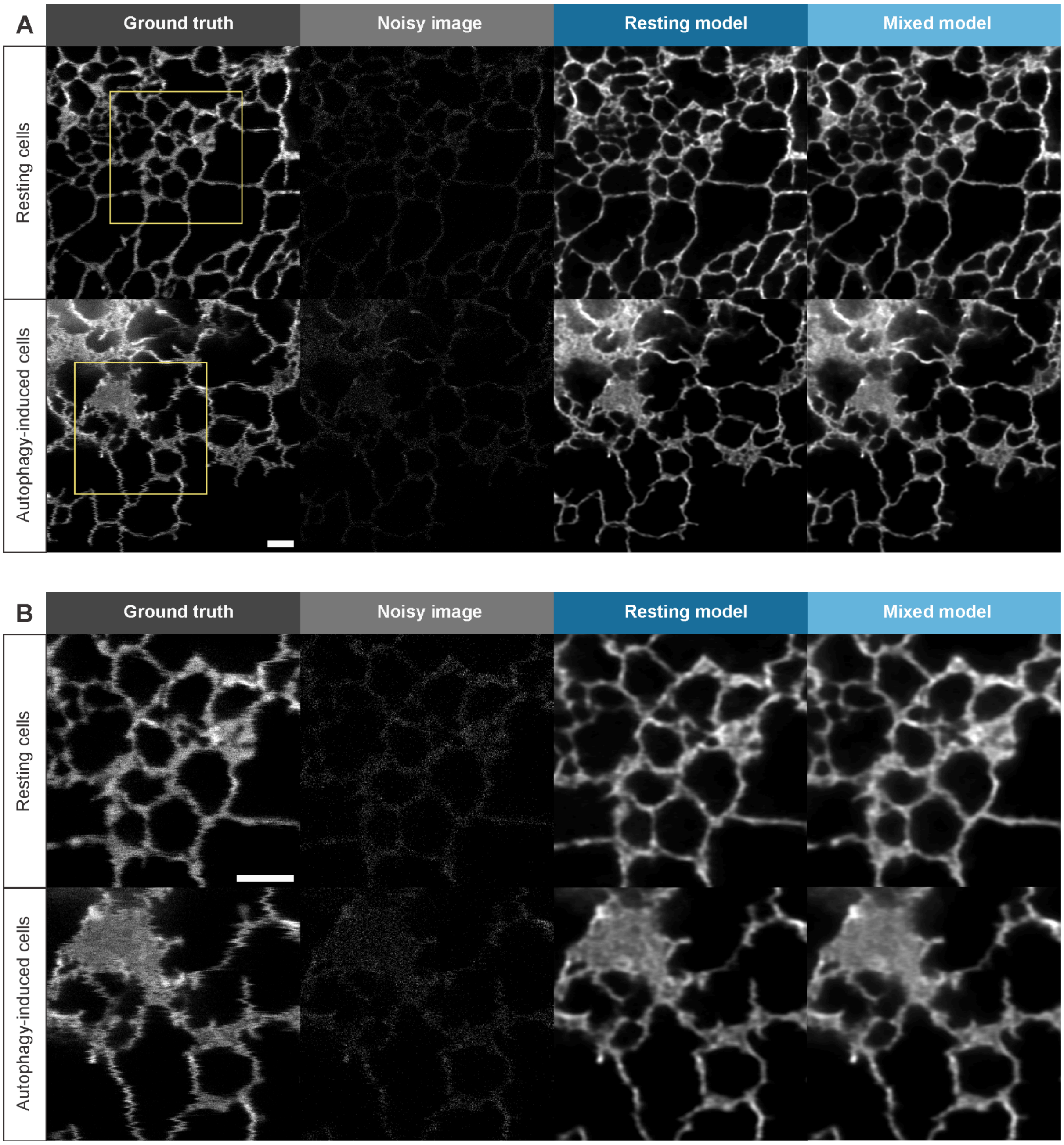

**SI Figure 6: Prediction output from different models. A** Representative ground truth and noisy images from cells in resting and autophagy induced state and their corresponding predictions using resting model and mixed model. Highlighted yellow box is the region of the zoom-in in **B**. Scale bars: 1 µm.

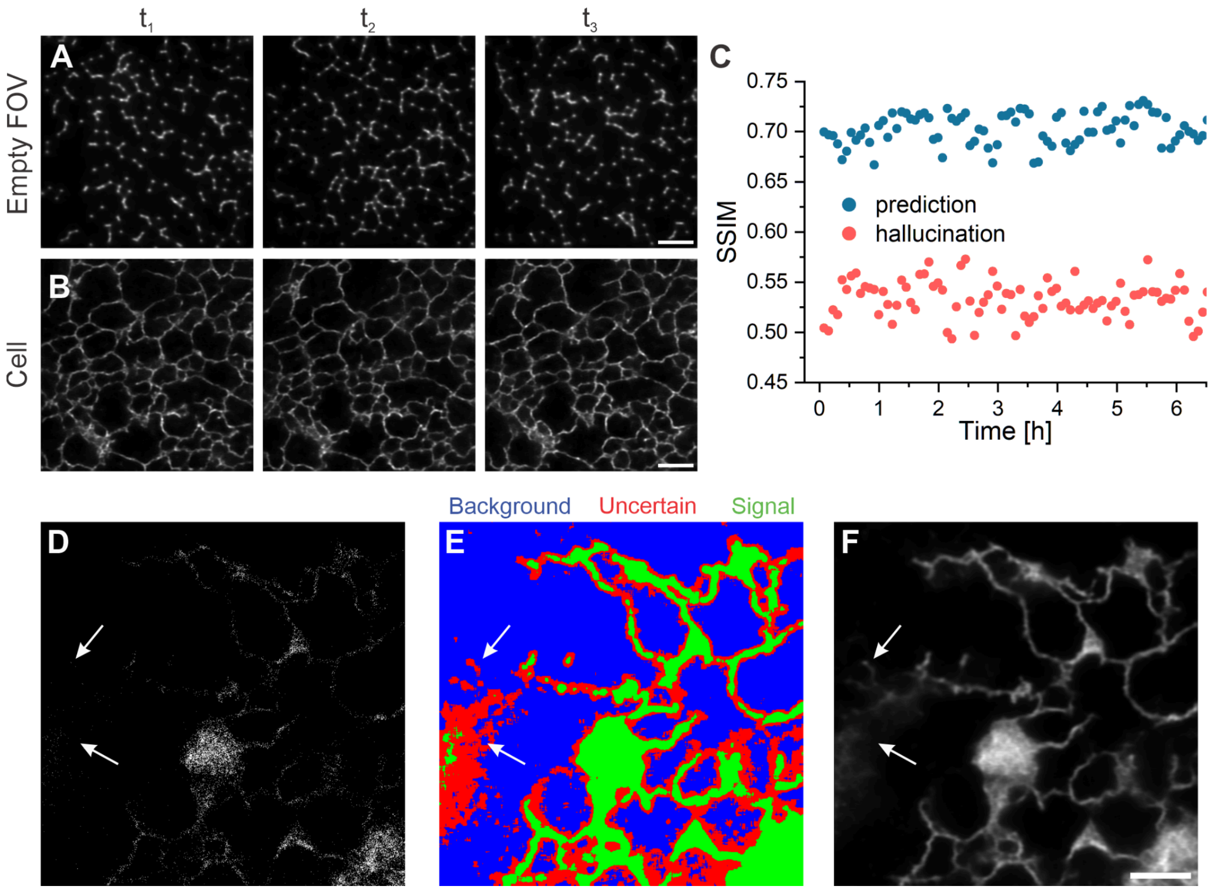

**SI Figure 7: Hallucination artifacts.** Predictions of a time series of background signals, which contain hallucinations. **B** Predicted images of a time series of ER in living cells. **C** To distinguish both cases, the SSIM for adjacent frames is calculated. The hallucinated structures appear randomly due to the random nature of noise and have a lower SSIM than a movie containing signal of interest. **D** Low-intensity input from treated movie condition. **E** Classified pixels as background, uncertain, and signal. **F** Predicted image, with white arrows indicating regions of sparse signal in the input data, which leads to hallucinations. Scale bar: 2 µm.

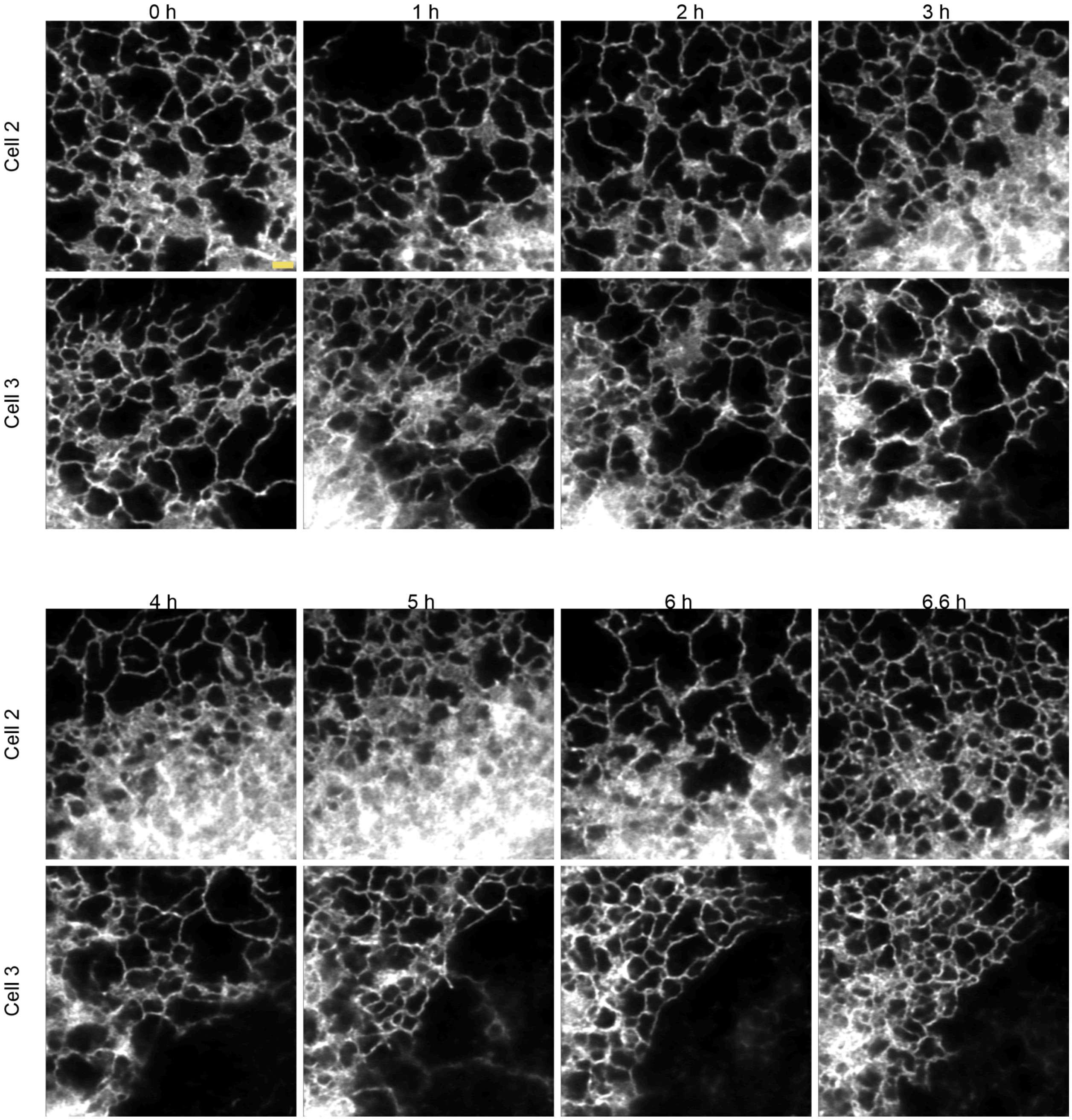

**SI Figure 8: Predictions of time series data of live-cell ER.** Representative predicted images of 2D planar low-intensity STED images of the ER at different time points acquired on two different live-cells (Cell 2: video S2; Cell 3: video S3). Scale bar: 1 µm.

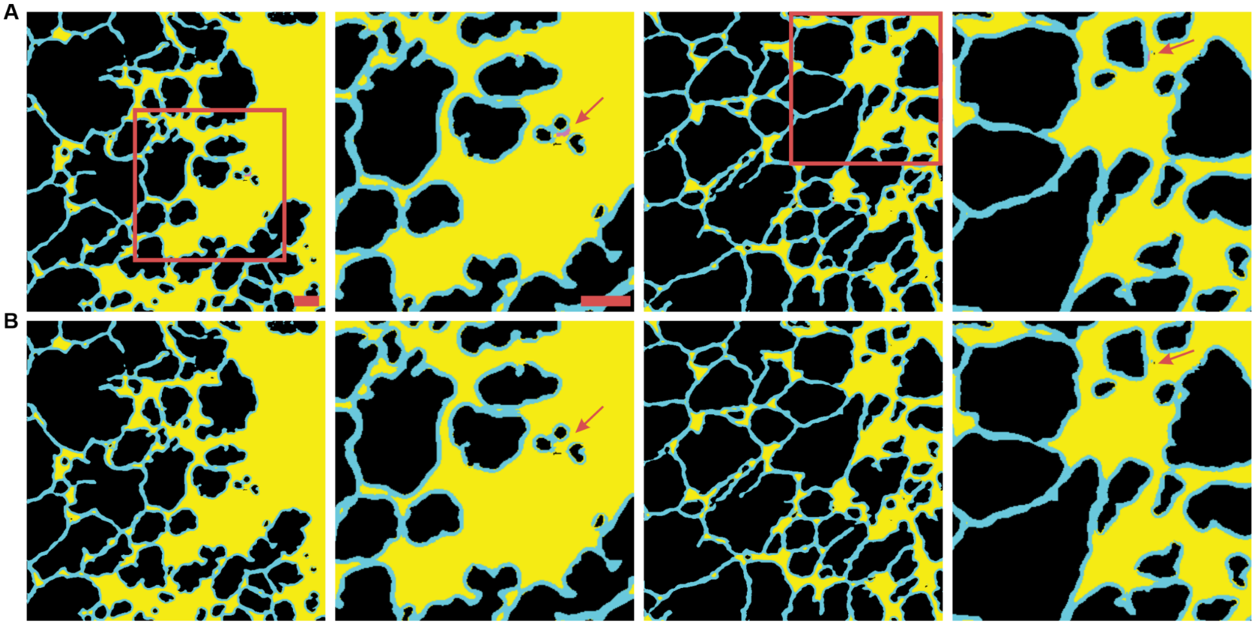

**SI Figure 9: ERnet segmentation output with and without sheet based tubes (SBTs)**. **A** Segmentation output of predicted ER images from ERnet visualizing tubes (cyan), sheets (yellow) and SBTs (pink) with corresponding inlays to the right. **B** The same segmentation output shown in **A** with SBTs merged with sheets (yellow). Scale bars: 1 μm.

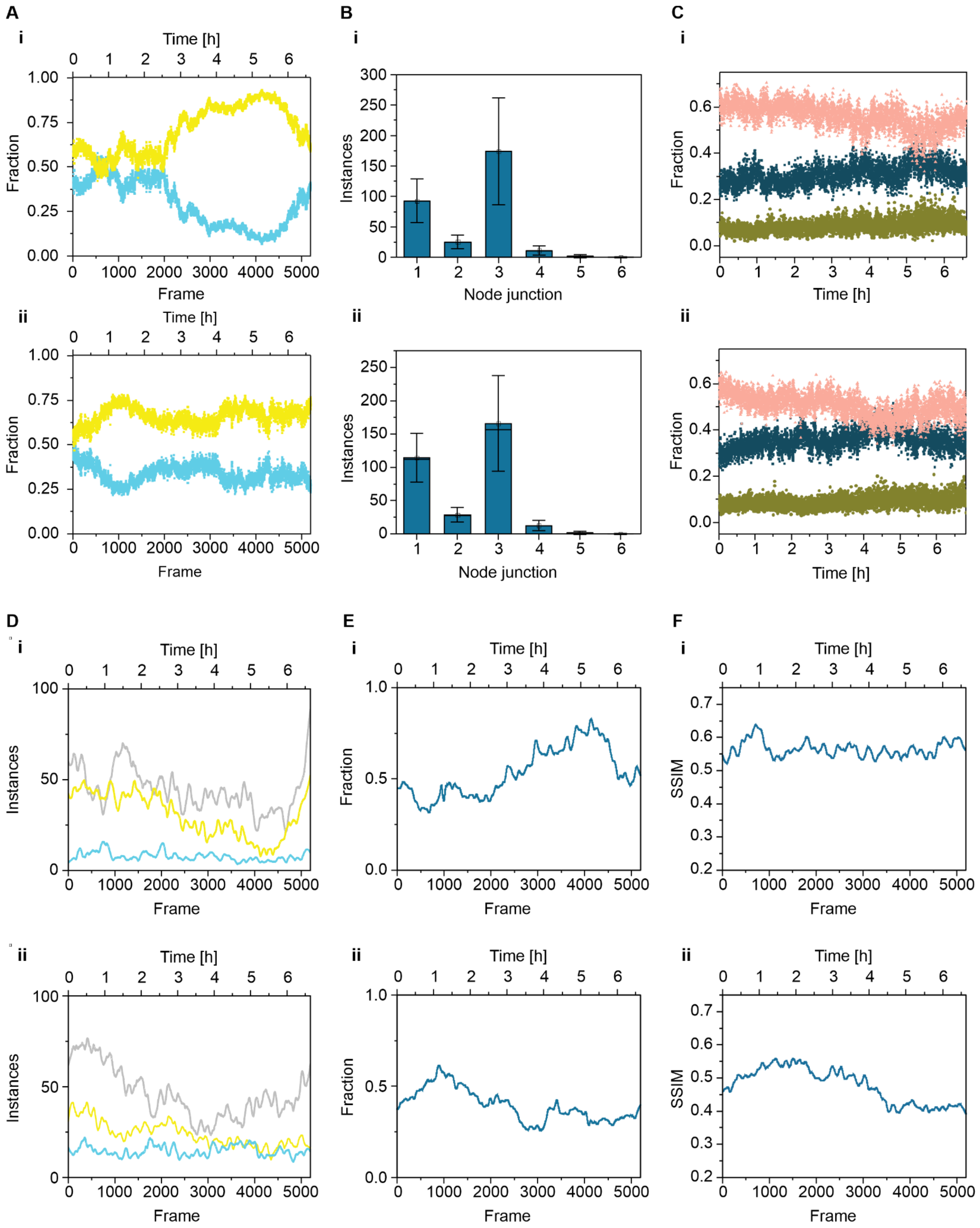

**SI Figure 10: Segmentation analysis of 2D time series of ER.** ERnet segmentation results on the predicted 2D planar time series of two different live-cells (**i** video S2; **ii** video S3) showing Tube (cyan) and sheet (yellow) fraction (**A**), the corresponding tube junctions averaged over all frames (**B**) and variations in one- (green), two- (blue) and three- (pink) way junctions over the whole measurement (**C**). D Instances of tubes (yellow), sheets (cyan) and gaps (light gray) varying over every frame for **i** and **ii**. **E** Fraction of foreground pixels and its variation over every frame for **i** and **ii**. **F** SSIM index of adjacent frames for all frames for the time series measurements **i** and **ii**.

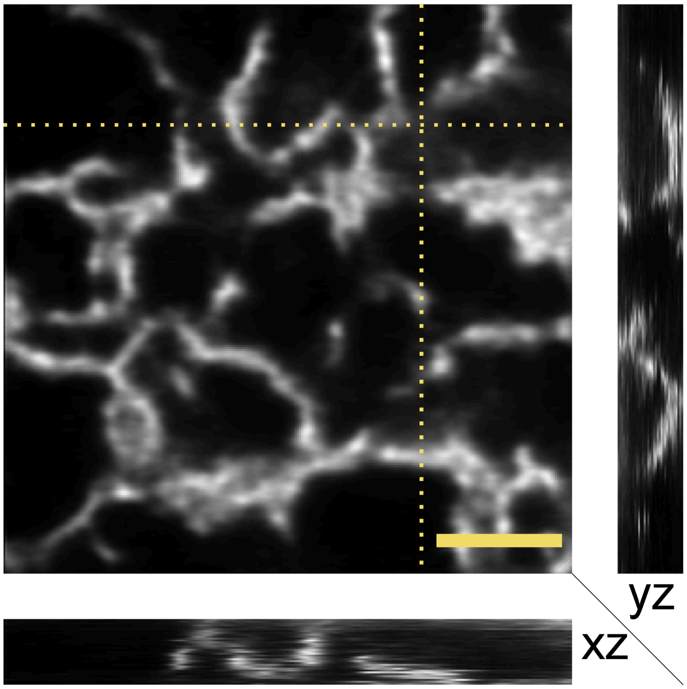

**SI Figure 11: Prediction of 3D stacks.** Prediction of the resting model for an input containing low-intensity volumetric STED images of the ER (calreticulin-KDEL). An exemplary xy plane is visualized and the yz and xz planes corresponding to the yellow dotted lines are shown on the left and bottom respectively. Scale bar: 1 μm.

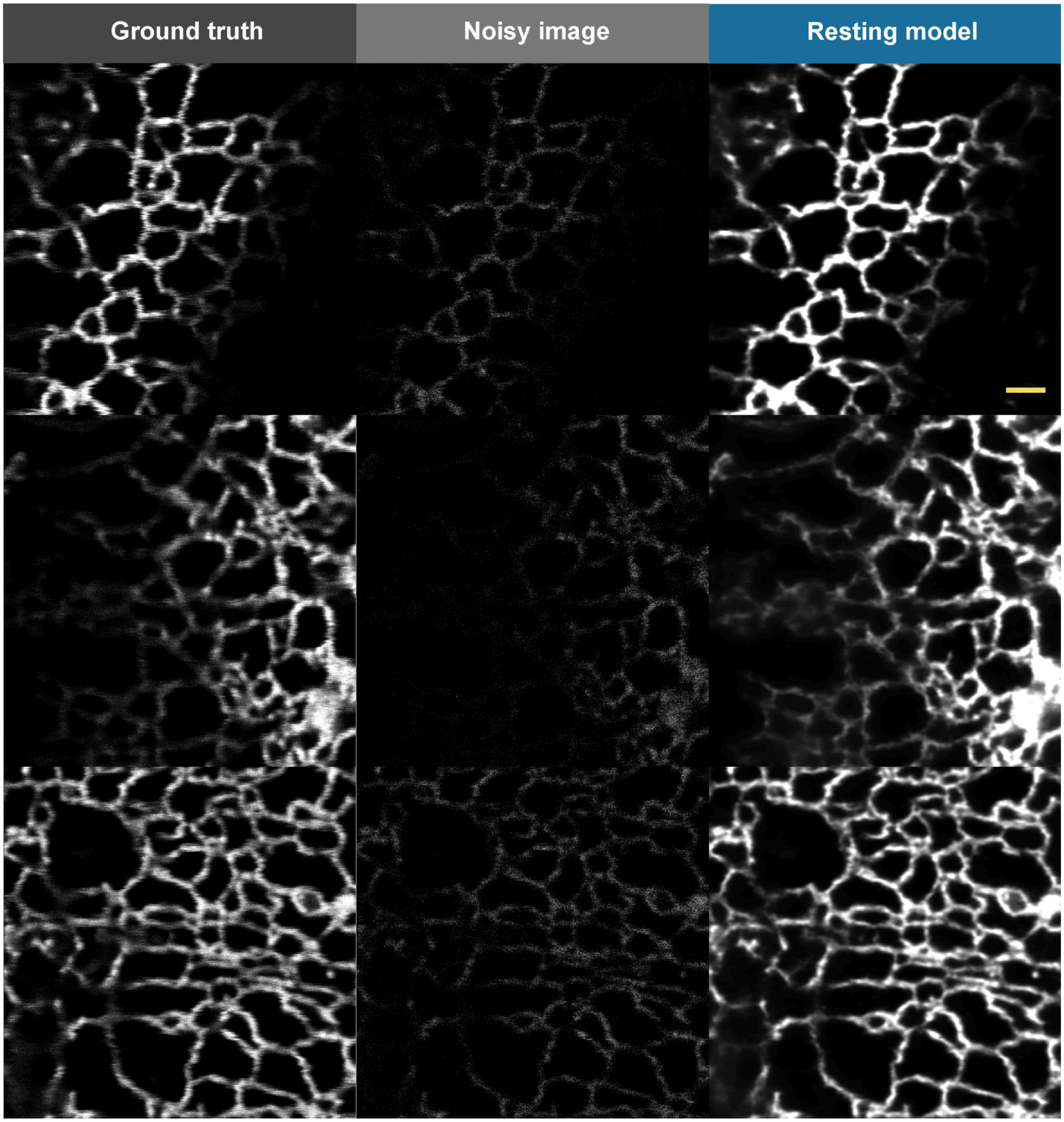

**SI Figure 12: Prediction of 3D controls.** Predicted output of planar images acquired with the same settings as the volumetric imaging including a top-hat PSF for the depletion laser beam. Scale bar: 1 µm.

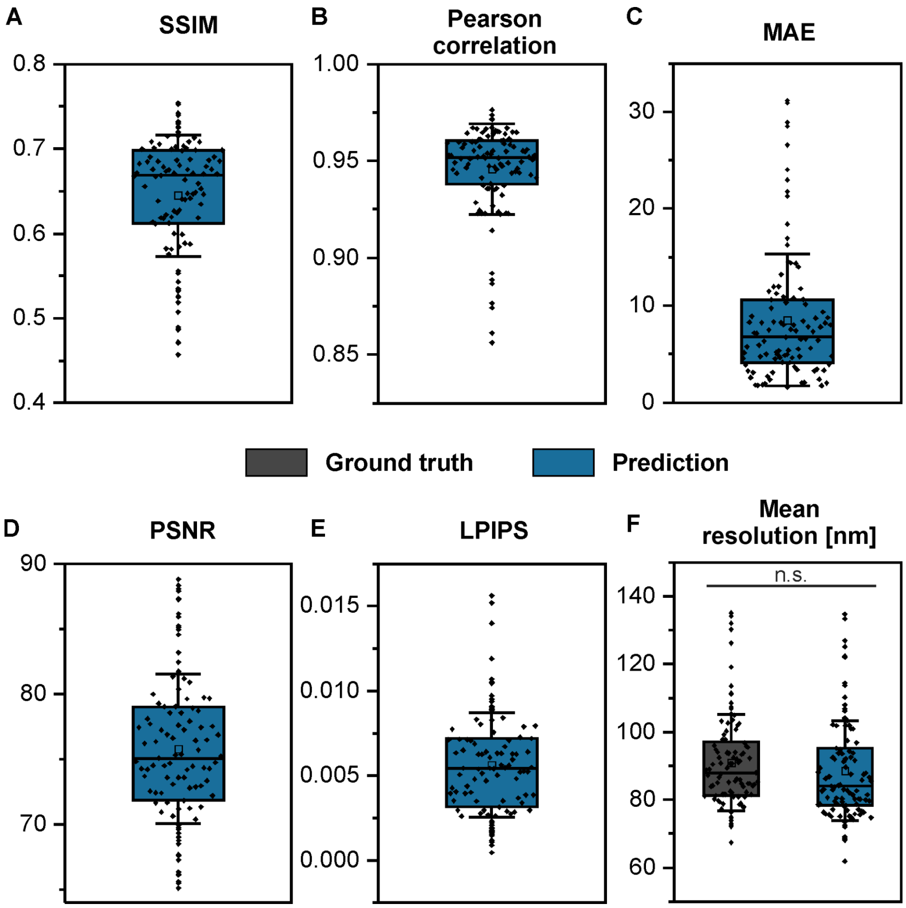

**SI Figure 13: Quality control metrics of 3D controls.** QC metrics of predictions from planar images acquired with the same settings as the volumetric imaging including a top-hat PSF for the depletion laser beam ranging from **A** SSIM **B** Pearson correlation **C** MAE **D** PSNR **E** LPIPS to **F** mean resolution from rFRC.

**SI Table 1: Hyperparameter importance of the resting condition.** The optimal parameter choices are highlighted in bold.

| learning rate | score | patch size | score | L2 regularization | score | activation | score | kernel initialization | score | clip value | score |
|---|---|---|---|---|---|---|---|---|---|---|---|
| 1E-4 | 13.00 | **304** | **28.28** | **0** | **35.39** | leaky relu | 48.79 | **Glorot uniform** | **45.25** | 0.1 | 32.23 |
| **1E-5** | **47.97** | 256 | 20.44 | 0.01 | 30.65 | **tanh** | **51.21** | Lecun uniform | 29.74 | 0.01 | 30.59 |
| 1E-6 | 39.03 | 200 | 25.75 | 0.001 | 33.96 | | | orthogonal | 25.01 | **0.001** | **37.18** |
| | | 128 | 25.52 | | | | | | | | |

**SI Table 2: Hyperparameter importance of the mixed condition.** The optimal parameter choices are highlighted in bold.

| learning rate | score | patch size | score | L2 regularization | score | activation | score | kernel initialization | score | clip value | score |
|---|---|---|---|---|---|---|---|---|---|---|---|
| 5E-4 | 4.56 | **304** | **25.95** | 0 | 32.73 | leaky relu | 47.93 | **Glorot uniform** | **37.51** | 0.1 | 32.69 |
| 1E-4 | 11.81 | 256 | 23.26 | **0.01** | **35.72** | tanh | 52.07 | Lecun uniform | 35.66 | **0.01** | **35.29** |
| 5E-5 | 28.29 | 200 | 24.98 | 0.001 | 31.54 | | | orthogonal | 26.83 | 0.001 | 32.02 |
| **1E-5** | **31.08** | 128 | 25.81 | | | | | | | | |
| 1E-6 | 24.27 | | | | | | | | | | |

**SI Table 3: Resting and mixed model parameters.** Unstated parameters are set as default values.

| **Resting model** | | | | | | | |
|---|---|---|---|---|---|---|---|
| lr | ps | bs | l2 | activation | kernel init | clip | epochs |
| 1E-5 | 304 | 6 | 0 | Leaky relu | Glorot uniform | 0.001 | 32 |
| **Mixed model** | | | | | | | |
| lr | ps | bs | l2 | activation | kernel init | clip | epochs |
| 1E-5 | 304 | 6 | 0.001 | Leaky- | lecun_uni- | 0.1 | 313 |

| | | | | | |
|---|---|---|---|---|---|
| | | | | relu | form |
